# A systematic review of professional society-backed biology education DEI research: The groups, research methods, and levels of analyses comprising the field

**DOI:** 10.1101/2024.09.19.613887

**Authors:** Candice Idlebird, Rebecca Campbell-Montalvo, Gary S. McDowell, Emily Blosser, Richard Harvey, Jana Marcette

**Affiliations:** Department of Social Sciences, Claflin University, Orangeburg SC, USA; Department of Curriculum and Instruction, University of Connecticut, Storrs CT, USA; Lightoller LLC, Chicago IL, USA; IGDORE, Chicago IL, USA; School of the Art Institute of Chicago, Chicago IL, USA; Department of Sociology, Anthropology, and Human Development & Family Science, University of Louisiana at Lafayette, Lafayette LA, USA; Department of Psychology, Saint Louis University, St. Louis MO, USA; Division of Academic Affairs, Montana State University Billings, Billings MT, USA

## Abstract

This integrative literature review analyzes the corpus of biology education research published in the main biology education journals of major professional societies. The goal of this analysis is to determine which approaches (including groups of focus, research methods, and settings/perspectives) from social science fields (i.e., psychology, sociology, and anthropology) are utilized in published peer-reviewed biology education research relating to diversity, equity and inclusion (DEI). Scoping how social science approaches are used in this area is important to understanding whether biology education research could benefit from complementary approaches that might advance praxis. This analysis found that research informing the biology education community draws heavily from psychological perspectives that are overwhelmingly not disaggregated (78% of articles identifying a group used a lumped together one), are by far more quantitative (58% used survey, 26% grades, 20% school data) than qualitative (17% used interview, 10% observation), and did not (72%) adopt structural approaches. The addition of missing contributions from social science is critical to advancing interventions to broaden STEM participation given that merging paradigms can offer more robust, multi-level explanations for observed phenomena. This has important implications for education, biology education, biology education research, social science, and research in related STEM fields.

## Introduction

Biology education research (BER) is a recently emerging field at the nexus of the natural and social sciences. In recent decades, BER has increasingly centered its work on investigating and framing student learning and engagement, with increased focus on promoting inclusion and equity in educational environments (Dirks, 2011). Much like in other STEM fields, retaining and engaging students in biology classrooms poses challenges, and this topic has taken on renewed importance in BER.

An incipient body of literature suggests BER favors quantitative methods, missing out on insights from qualitative and mixed methods approaches (e.g., Lo et al., 2019). While using quantitative methods provides the advantage of statistical generalizability, there are well known limitations to their sole use, particularly in understanding experiences of marginalized groups, for reasons including sample size and foundational design (Slaton & Pawley, 2018; Garcia et al., 2018). To address sample size issues as well as due to misunderstandings about social theorizing, quantitative approaches can often aggregate all marginalized groups together (e.g., including Black, Latinx, and American Indian in one group) as well as fail to see within group differences (e.g., African American, African). At the same time, much educational research relies on individual level approaches to student success, rather than those that consider structural or systemic issues underpinning outcomes.

Given the importance of retaining students and DEI-focused work on broadening participation in the field, this article contributes to the literature by assessing the scope of studies in BER. Specifically, we take a first step toward this goal by investigating and analyzing the current state of the field of BER that focuses on DEI. To do this, we analyze papers published in four top biology journals overseen by the major biology/biology education professional societies. We focus on publications by professional societies (e.g., the American Society for Cell Biology, American Society for Microbiology, National Association of Biology Teachers), given that practitioners often look to them for information and professional development when it comes to their own pedagogies and educational research efforts. Indeed, societies’ journals directly inform the research and teaching of practitioners, including such journals as *CBE—Life Sciences Education* or SABER conference papers (Lo et al., 2019).

More broadly, scientific societies impact disciplinary culture and norms both through their defining the disciplinary field, and as influencers of a range of stakeholders with agency to make changes individually (Brownell & Tanner, 2012; Greenwood et al., 2002; Leibnitz et al., 2021). Scientific societies have demonstrated an interest in promoting diversity, equity, and inclusion within STEM environments with increasing interest from biology-related disciplines in particular (Abernethy et al., 2020; Campbell-Montalvo et al., 2020, 2022a; Leibnitz et al., 2021; Mourad et al., 2018; Segarra et al., 2020a; Segarra et al., 2020b; Wilton et al., 2022). We add that when it comes to consideration about what exactly gets published in BER journals, Dolan (2012) brings forward the tension between the desire of researchers to publish academic research (perceived as being more rigorous and a well-known expectation for tenure) over publishing the implementable classroom tools sought by practitioners reading the journal.

## Background

Recent scholarship in BER provides some insight into the current state of the field, suggesting a biology education research tradition mostly excluding qualitative, disaggregated, and structural approaches. Yet this existing research is somewhat dated and therefore might not reflect a present likely more scaffolded upon increased emphasis in DEI in STEM more broadly. For example, in one of the few articles which has undertaken a similar inquiry as ours, Lo and colleagues performed a content analysis of peer-reviewed research articles in biology education research (i.e., in *CBE—Life Sciences Education*). They focused on research articles between 2002 and 2015, and also included conference abstracts for the Society for the Advancement of Biology Education Research (SABER) between 2011 and 2015. The authors concluded that these articles and abstracts relied heavily on causal research questions and quantitative methods (Lo et al., 2019). In addition, the authors found an important distinction among SABER abstracts: research talks were more likely to use quantitative methods, and posters were more likely to report qualitative methods, with poster presentations being perceived to have lower prestige. A crucial point Lo and colleagues made in interpreting these findings is that most of the research that was being carried out on and in biology classrooms at that time was by practitioners who had “crossed-over” to biology education research from the biological sciences where they had received training in causal, quantitative research methods, potentially without benefitting from training on social science methods and theories (Lo et al., 2019).

Lo and colleagues also found that the setting of biology education research draws more on a classroom context (Lo et al., 2019). On one hand, studies of this nature may focus on using discrete metrics, such as grades or academic performance indicators, for understanding student success. On the other hand, fields such as sociology and anthropology draw on qualitative methods to better understand the in-depth perspectives of individuals, and capture important elements at the university, cultural, and society context in understanding social forces at play in STEM education. Using narratives made possible through such methods as interviews, ethnography, and focus groups reveals insights not capturable in grades and other data (Aldridge & McChesney, 2018; Bandura, 1977; Idsoe, 2016; Mishra, 2020; Tinto, 2006).

Qualitative approaches help researchers understand how and why people experience what they do. For instance, Bucholtz et al. analyzed ethnographic data from a study with undergraduate math and science majors that yielded 439 hours of audio video content of interactions amongst 118 participants who were part of a larger pool (n=496, Bucholtz et al., 2012). Interviews and a survey offered additional data for triangulation and to complement the strengths of the other methods. The authors took a close look at how one lab group interacted during their weekly meetings in an honors first-year chemistry class over the course’s 10 weeks. There were one man and two women Chemistry majors in the group (the participants’ race was not included by the researchers). The authors articulated how the team’s dynamic shifted during this time as the two women first acknowledged the man as a competent team member but then later imposed “a socially and academically marginalized persona through habitual mockery and criticism” even though his grades were similar, though sometimes slightly lower than theirs (Bucholtz et al., 2012, p159). In the end, the authors argue that understanding how student identities unfold in labs and classrooms requires attention to how time unfolds in interactions, and the underlying ideology which links meaningful resources across contexts. Such focus on ideology and time when considering identity is a key consideration in sociology and anthropology, rather than psychology (see Rosa & Flores, 2017 for example).

Relatedly, in scientific research, White male participants have been used as the primary population studied, and results were then generalized to all, including marginalized groups (Lo et al., 2019). Further, research is often carried out under the White gaze (Paris, 2018), lumping together and naming marginalized groups as deficient, reproducing settler logics. For instance, work can aggregate Black/African American, American Indian, Asian, Latinx, and other Persons Excluded due to Ethnicity or Race (PEER, Asai, 2020) into one group. To help address this, NIH has supported a requirement to include racial/ethnic data in studies since 1993 (National Institutes of Health, 2017). The requirements state that data on race, ethnicity, and gender should be reported, and participant populations should be diverse. However, African American and Latinx people account for less than 10% of study participants (Pérez-Stable, 2018). Additionally, gender and sexual marginalized groups (e.g., LGBTQIA+) are regularly excluded from research studies (Freeman, 2020), but when included can be lumped together in one group despite the vastly different impacts that having gender marginalized (transgender, non-binary) versus sexual marginalized (lesbian, gay, bisexual) identities has (Campbell-Montalvo et al., in press). The invisibility of whole communities of participants can lead to inaccuracy in attempts to make theoretical assumptions about how research impacts communities, particularly a problem in attempts to parse out how Asian students (e.g., underrepresented Southeast Asian students, e.g. Cambodian, in comparison to East Asian students, e.g., Chinese) experience STEM (Her, 2019; McGee, Thakore, & LaBlance, 2017). Better inclusion of a diversity of groups is required in order to foster a BER paradigm that can better inform approaches to success (Freeman, 2020; Intemann, 2009; Lo et al., 2019; Pérez-Stable, 2018; Yonas et al., 2020).

At the same time as issues of methodological emphasis and group operationalization are germane, the level of analysis used in BER is also a paramount concern. Level of analysis relates to how students’ struggles are framed and whether individual versus structural frameworks are used to understand student experiences. A large body STEM education research mobilizes individual level approaches rooted in psychology to diagnose problems such as poor outcomes, grades, and student attrition, without appropriately overlaying the structural conditions underpinning these issues. Such an approach can back classroom interventions (e.g., Lo et al., 2019) assuming small tweaks to student experiences can singularly drive success, without understanding how classroom contexts and pedagogies intersect and are affected by larger systemic issues.

Sociological and anthropological approaches often center the group and structural level. For example, Campbell-Montalvo et al. draw from sociology as they mobilize social network theory (Lin, 2017) to understand how people in a student’s social network impact their persistence in their engineering major (Campbell-Montalvo et al., 2022b; Campbell-Montalvo et al., 2022c). First, drawing from interviews with 55 women and underrepresented students, the researchers found that students experienced differing engineering department climates because of how others reacted to their identities (Campbell-Montalvo et al., 2022a). For example, Black students experienced more stereotype threat (a concept born in social psychology (see Steele & Aronson, 1995), a point the wider literature has already identified as anti-Blackness of STEM (Bullock, 2017; Cedillo, 2018; Martin et al., 2019; Nxumalo & Gitari, 2021; Vakil & Ayers, 2019). This resulted in such students having to not only be aware of how others likely perceived them, but to also make sophisticated calculations about how they should behave in order to manage others’ impression of them (a concept known as stereotype management, see McGee & Martin, 2011; McGee, 2016). Yet, having people in an undergraduate engineering student’s social network helped them persist in their degree program--a pattern suggested in this qualitative research and confirmed through inferential statistics on surveys with over 2,000 undergraduates (Smith et al., 2021).

In Campbell-Montalvo et al., the mechanisms at play wherein social networks impacted persistence came from individuals who saw students as total people, individuals who were close to students like their parents, relatives, or teachers who were often engineers or scientists (Campbell-Montalvo et al., 2022b). Women and Black and Latinx students received advice from these individuals that they should expect to be stereotyped, and when that happened, the advice served as an armor to help students make sense of what happened in a way that did not harm their fit in engineering and they were able to persist (Campbell-Montalvo et al., 2022a). Drawing from anthropology, the authors centered cultural model theory as they theorized about students’ cultural models of engineering success (Smith et al., 2015). Students’ cultural models of engineering success are the ways that students understand and think about how to be successful in engineering. One component of this type of cultural model is the notion of sense of belonging or fit, wherein fit is a *structural* outcome based upon the social and academic environment the student experiences in their engineering department based largely on department, university, and societal policies, histories, and forces (Campbell-Montalvo et al., 2022).

Similarly, the researchers argued that students participating in professional engineering societies were more likely to persist than those who did not because of the common social identities of participants and those in the societies (Campbell-Montalvo et al., 2022c). That is to say, Black students participating in the National Society of Black Engineers reported that being around other engineers like themselves, seeing them succeed and receiving advice from them helped them maintain their fit in engineering. Thus, the mechanism at play is a range of advice received from people in a students’ social network that had much to do with those people sharing similar characteristics with the students. In these studies (Campbell-Montalvo et al., 2022b; Campbell-Montalvo et al., 2022c; Smith et al., 2021), the power in the advice comes from people with social characteristics like the students is known as alter homophily--a concept the authors borrow from sociology (McPherson et al., 2001). Ultimately, given the anti-blackness of STEM, the differential treatment of Black students in engineering compared to Latinx and other counterparts, the authors argue that social networks and their resultant effects on fit are an important factor to consider when designing interventions to address STEM inequities; yet they caution that approaches aimed at fixing such students must also exist with approaches that seek to improve the STEM climate in which marginalized students are learning (Campbell-Montalvo et al., 2022a). Additionally, these findings explained here relating to nuanced identity implications in which Black students had different experiences from Latinx students were made possible by the team’s disaggregated approach. Yet more work needs to be done to further investigate intergroup differences, including those among Black, Asian, Latinx, and other students. In any event, these handful of exemplar articles illustrate the power of work on STEM inclusion when multiple research methods, disaggregated grouping, and numerous levels of analyses are deployed to robustly measure and deconstruct phenomena to name processes to inform application.

Going back to the state of BER when it comes to methods, group aggregation, and analytical level, we note that, given these characteristics of BER found in work reviewed almost about 10-20 years ago in Lo and colleagues (2019), there have been calls for expanded approaches in BER. These calls have asked for the inclusion of Discipline Based Educational Research (DBER), the incorporation of social science fields and approaches, and the linkage between researchers and practitioners. For example, efforts to integrate with other DBER communities include expanding relationships with the long-established field of physics education to address STEM teaching and learning as well as inclusion and institutional change (Shipley et al., 2017). Another example of calls for expansion are seen in Dolan’s (2012) perspective piece, which reviewed the literature comprising discussions in BER-focused journals and national reports echoing calls for greater collaboration and communication between BER researchers and social sciences researchers, and with biology classroom practitioners.

In the same vein of collaboration, Dolan’s later (2015) editorial in *CSE—Life Sciences Education* adds to the case to explore the utility of other fields, particularly in the social sciences, to incorporate new theories and methods into BER, which would allow researchers to explore important consequential research questions and outcomes (e.g., how biology students develop science identity and a sense of belonging in STEM (Dolan, 2015). Moreover, Peffer and Renken (2016) suggest incorporating natural science traditions into DBER to provide a better and more robust understanding for science collaborative research to improve DBER. And when it specifically comes to BER, Akman and colleagues (2020) highlighted the efforts of four education reform organizations (BioQUEST Curriculum Consortium, MathBench Biology Modules, QUBES [Quantitative Undergraduate Biology Education and Synthesis], and the Intercollegiate Biomathematics Alliance) to effect systemic change in the mathematics biology education community (Akman et al., 2020).

Ultimately, it remains to be seen as to how these interdisciplinary calls have actually impacted BER, particularly BER related to DEI. Thus, understanding the scope of more current DEI-focused BER would illuminate the gaps and offer important insight to augment BER with additive approaches. Therefore, the following research questions guide this study:

1. What types of methodologies are used in biology education research?
2. How are marginalized groups defined and conceptualized in biology education research?
3. What types of frameworks do biology education researchers use to draw conclusions (with a focus on individual versus structural frameworks)?

Knowing the answers to these questions, and therefore, how far BER has come, will help inform more expansive approaches in the field, approaches seen in publications outside of biology societies where researchers have utilized a range of approaches to better understand and address STEM undergraduate inequity (Cech & Blair-Loy, 2019; Cech & Waidzunas, 2021; Corwin et al., 2020; Leyva, 2016; Leyva et al., 2021, 2022; McGee & Martin, 2011; McGee, 2016; Morton et al., 2020; Morton & Parsons, 2018; Secules et al., 2018, 2021; Smith et al., 2014).

## Methods

In order to explore themes addressed above, we use an integrative approach. We collected a key sample of research articles on the topic of diversity, equity, and inclusion in biology education research, and undertook data analysis of these articles to identify trends in this body of scholarship. We selected articles from journals associated with national biology professional societies, including such journals based upon the immense role of national professional societies in fostering community within STEM disciplines. These education articles further have a wide readership among biology instructors who are not themselves education researchers. These journals are discipline-based, where articles are written by higher education biology practitioners themselves for their community, and therefore are illustrative of a community of thought among biology practitioners.

To address our research questions, we collected all abstracts from research articles from five professional society-run biology education research journals, including all articles spanning January 2017 through July 2020 (Table 1). The societies include the Association of College and University Biology Educators, American Society for Biochemistry and Molecular Biology, American Society for Cell Biology, American Society for Microbiology, and National Association of Biology Teachers. Collectively, their membership includes students, researchers, educators, and bioscience professionals across various biological science disciplines. Their journals include *Bioscene: Journal of College Biology Teaching, Biochemistry and Molecular Biology Education, CBE Life Sciences Education, Journal of Microbiology and Biology Education,* and *The American Biology Teacher.* These journals’ foci and scopes are specifically geared towards biology education (Table 2).

**Table 1:** Journals Identified for the study

| **Biology Professional Society** | **Membership defined by society** | **Acronym** | **Professional Society Journal** | **Journal Acronym/Abbreviation** |
| --- | --- | --- | --- | --- |
| Association of College and University Biology Educators | “ACUBE's members are a group of post-secondary educators dedicated to excellence in teaching biology.”<br>http://www.acube.org/index.php | ACUBE | Bioscene: Journal of College Biology Teaching | Bioscene |
| American Society for Biochemistry and Molecular Biology | “biochemists and molecular biologists”<br>https://www.asbmb.org/membership | ASBMB | Biochemistry and Molecular Biology Education | BMBE |
| American Society for Cell Biology | “Research scientists in academia and industry; Postdoctoral fellows; Educators (community college instructors); Graduate students; Undergraduate students...scientists who study the biology of the cell”<br>https://www.ascb.org/membership/ | ASCB | CBE Life Sciences Education | CBE-LSE |
| American Society for Microbiology | “researchers, educators and health professionals”<br>https://asm.org/about-asm | ASM | Journal of Microbiology and Biology Education | JMBE |
| National Association of Biology Teachers | “teachers from all settings, from K-12 to postgraduate to informal”<br>https://nabt.org/MembersArea | NABT | The American Biology Teacher | ABT |

**Table 2.**
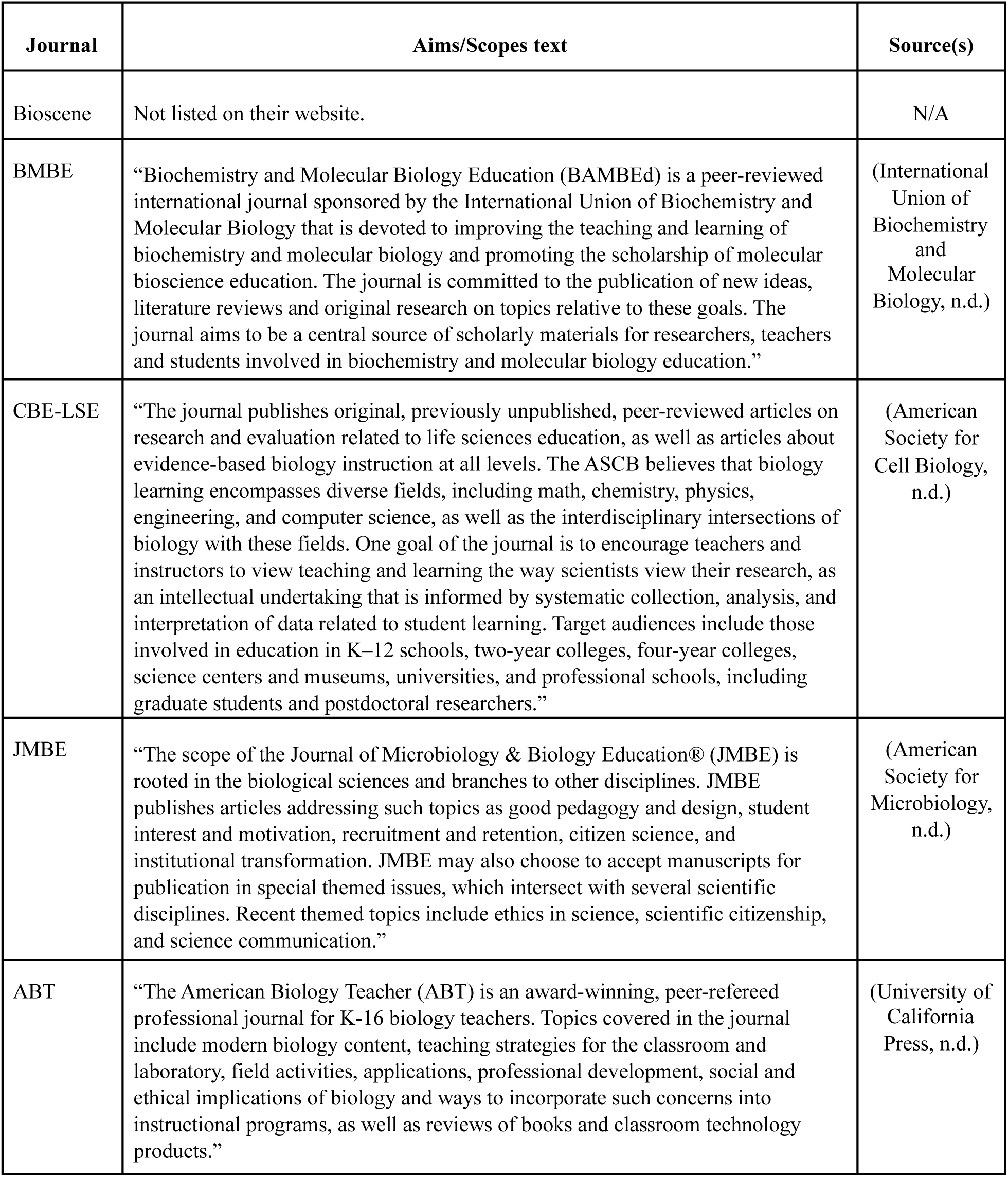
: Aims and scopes of journals

We note that the sample does include some special issues with calls for papers on topics of diversity, equity, and inclusion. The 3.5-year window (January 2017 through July 2020) of inclusion spans the most recent articles prior to our team’s embarkment on analysis, provides a large but feasible amount of work to analyze, and complements existing analyses on periods prior to this point.

As part of the initial review, we sought to estimate the number of research articles that might comprise the sample for close analysis. To do this we first began with defining search terms for articles of interest within the journal sample set. Two co-authors who are subject matter experts with experience in communities of biology education practitioners and an interest in inclusive science practices, met several times to discuss how inclusion is conceptualized at meetings of biologists and biology education practitioners. From these discussions, the co-authors met with the research team (which included additional subject matter experts from complementary fields such as social science) to discuss the search terms. Based upon this, the two co-authors conducted a search of articles in the journals identified using the keywords “diversity,” “equity,” and “inclusion.” Articles appearing with these three search terms were reviewed and the keywords associated with the articles were discussed as a team to create the final list of search terms: “broadening participation,” “underrepresented,” “under-represented,” “disadvantaged,” “diversity,” “equity,” “inclusion,” “anti-racist,” “minority,” “disadvantaged,” “pell-eligible,” and “PELL eligible.” We counted the approximate number of all articles associated with these terms and estimated the total number of articles available for analysis to be 1,293 (Table 3).

**Table 3:**
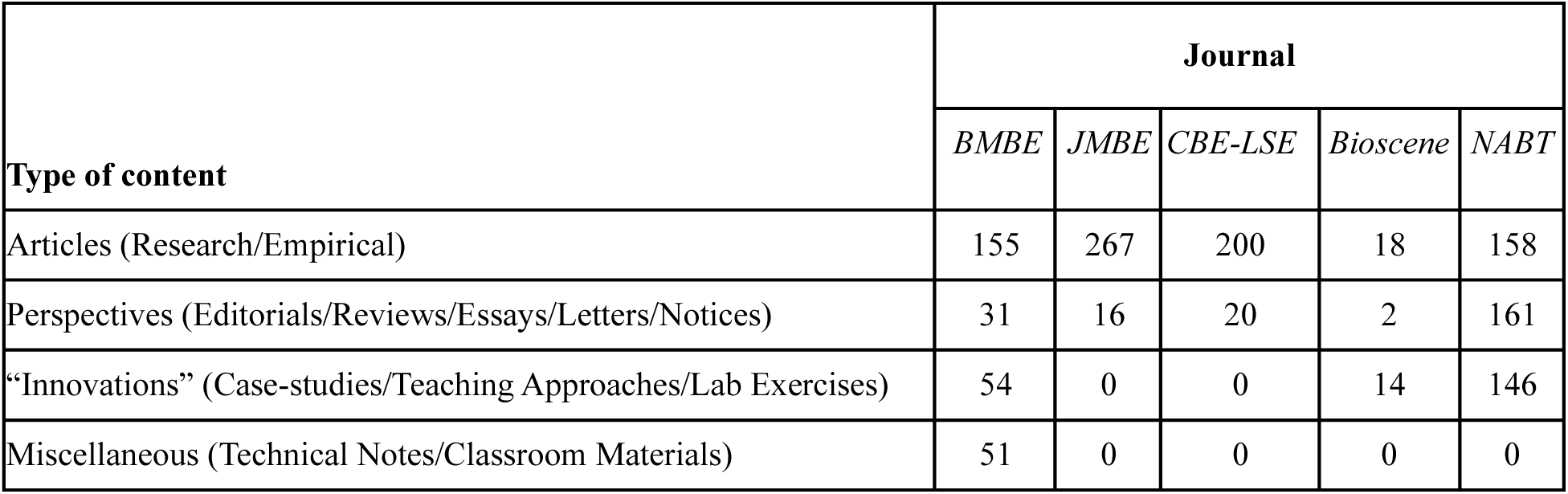
Estimation of articles published in five National Professional Society Biology Education Journals from 2017-2020 on diversity, equity, inclusion (2017-July 2020)

One team member then parsed these 1,293 articles out by type (i.e., research, perspective, innovations), shown in Table 3. This parsing was done based upon review of the journal’s Table of Contents, showing that the sample consisted of 798 research articles, 230 perspectives, and 214 innovations. There were also 51 miscellaneous articles in *BMBE*, which consisted of technical notes or specific classroom materials that could be used to teach biology. We note that *JMBE* and *Bioscene* include conference abstracts, which were not analyzed as part of this work. *First Round of Coding: Confirmation of Research Study and Underrepresented Group of Focus* In the first round of coding, we sought to confirm that our set of 798 articles indicated as research articles in the journals’ Table of Contents were actually empirically-based studies. We also wished to establish the categories of underrepresented groups on which they focused. This was done through systematic coding by four coders using a codebook with two codes that included: 1) Abstract is a research study; and 2) Study is about any underrepresented groups. In the codebook, underrepresented groups were defined according to the National Science Foundation operationalization (National Science Foundation, 2021) including underrepresented racial/ethnic marginalized groups (URM), women, persons with disabilities, and people with sexual and gender marginalized identities, but we also counted any other groups underrepresented groups mentioned as such.

During this first round of coding, our team included four coders (one biomedicine Ph.D.-holding interdisciplinary researcher, two HBCU undergraduate students—one majoring in sociology and one majoring in biology, and one graduate psychology student at a PWI) who met bi-weekly to discuss any coding that lacked agreement of two or more coders. There were also two additional subject matter experts (a sociology PhD-holding HBCU assistant professor, and either of: a biology PhD-holding former HBCU assistant professor who transitioned to an administrative role at a PWI; or a psychology PhD professor at a PWI) at the meetings to provide more insight and assist with agreement on coding. The specific codebook used during this time is shown below.

<u>Codebook 1:</u>

Is this a research study?

1. Yes
2. No

Is the article about an underrepresented group?

3. Yes
4. No

The inclusion criteria we used in the initial review and first round of coding were that the study was published in an education journal listed in Table 1; that the study was a peer-reviewed article or report with empirical findings (i.e., conclusions from quantitative or qualitative data were included); that the study was published between January 2017 and July 2020; and that the study explored diversity, equity, and inclusion in the context of biology education. Articles were excluded if they related solely to the study of a biological process. Using these methods, we coded the 798 abstracts reported by the journals’ Table of Contents as research articles, and we determined 645 to be research studies based on review of the full abstracts. Of the 645 research studies that appeared with our initial search terms, 93 were articles focusing on an underrepresented group.

To determine coders’ reliability with our approach, we calculated intercoder reliability across the four coders on the first two codes on 5% of the initial articles (n=5). For the first code on whether the article was a research study, the four coders applied the same code across all coders 87% of the time. For the second code, on whether the article was about an underrepresented group, the four coders applied the same code 97% of the time.

In the second round of coding, and in concert with our research questions, we identified the specific demographics discussed within each article, identified the research methods used, and considered the level of analysis. This second round of coding was done on the 93 articles that were coded in round one as being research articles about diversity, equity, and inclusion involving an underrepresented group. This round involved three of the four coders, with the undergraduate biology student not participating in this round. These three coders coded each set of abstracts for the second round of coding. We divided the abstracts into three groups of 31 and utilized Codebook 2 on one group of abstracts at a time. After each individual coded a group of 31, the coders and wider research team met to discuss any coding discrepancies in that group and arrive at consensus in cases where less than two coders were in agreement. The specific codebook used during this time is shown below.

<u>Codebook 2:</u>

Race/Ethnicity (select all that apply)

1. African American
2. American Indian
3. Asian
4. Latinx
5. White
6. URM as one group
7. Other racial/ethnic grouping

Sexuality/Gender identities (select all that apply)

8. Sexual minority group (Lesbian, Gay, Bisexual, Queer)
9. Gender minority group (Transgender, non-binary)

Level used (choose one)

10. Individual level (e.g., motivation)
11. Classroom level (e.g., teaching strategies)
12. Structural (e.g., university diversity programs)
13. Other/unsure

Research methods used (select all that apply):

14. Survey
15. Observation
16. Interview
17. Student grades
18. School data (i.e., enrollment/graduation rates)
19. Review of literature
20. Other

Prior to deploying the first codebook, the entire main research team of 8 met over the course of several months to construct the full codebooks (one and two) based upon the project’s research questions, previous literature, and initial review. These meetings usually included the 4 coders and 4 researchers involved in this project. Right before using the first codebook, the team used multiple meetings to code several samples as a group. During this process, the team discussed the meaning of each code, when to use them, and also refined the codebook to be usable. The four coders were then given a sample set of 40 abstracts to code, and the team met to review inconsistencies and learn how to code. During the initial review as well as the two rounds of coding, coders were trained on the codebooks and agreed-upon norms of deploying it, such as erring on the side of caution—if they were not sure whether an article should be counted as fulfilling a certain criterion they should code it as a ‘yes’ and discrepancies would be further discussed during team meetings. The training included discussion of how to capture data within a code, for instance, the code of Latinx in the second codebook was to be deployed in instances where Latino, Latinx, Hispanic and similar terms were used—though we acknowledge the terms are not synonymous. To be clear, the 93 articles were not read in full, and this study included only analysis of the abstracts.

## Results

The following is a description of the 93 abstracts about research articles relating to diversity, equity, and inclusion that were coded by the team. Though the initial sample drew more broadly from the range of journals, due to how the abstracts fit the criteria, we ended up closely reviewing a larger proportion of articles from some journals (namely, *CBE Life Sciences Education* [n=70] and then *Journal of Microbiology and Biology Education* [n=15]) than others (*Biochemistry and Molecular Biology Education* [n=4], *Journal of College Biology Training* [n=4]), given that our review showed the former pair of journals had mores abstracts than the latter that described research articles according to our operationalization described earlier.

In looking at the articles we analyzed, 60 of the 93 articles specified a focus on any race/ethnicity group in the abstract. For the 60 articles that did mention race/ethnicity, most (n=53) of these articles named one specific racial/ethnic grouping of focus: most commonly a grouped URM or similar designation (n=47), followed by a focus on African American/Black (n=3), Latinx (n=2), and then Asian (n=1). The seven remaining abstracts emphasized more than one specific race/ethnic grouping of focus, with three focusing on Black/African American and Latinx and two on Black/African American, Latinx, and American Indian, and the final two had two additional group combinations. In terms of meaning, arguably, papers using a URM denotation and those grouping together Black/African American and Latinx and Black/African American, Latinx, and American Indian have similar study foci.

Looking at all of the 60 abstracts together, noting that seven focused on more than one grouping (i.e., Latinx and Black, both), the URM category was emphasized in 47 (78%) of the abstracts that focused on a group, followed by abstracts centering on African American/Black participants (n=9; 15%) and Latinx participants (n=8; 13%). American Indian, other racial/ethnic grouping, and Asian were specifically mentioned in two to three abstracts each. Table 4 depicts the list of the racial/ethnic categories used in the articles, along with how many abstracts each was found in.

**Table 4.**
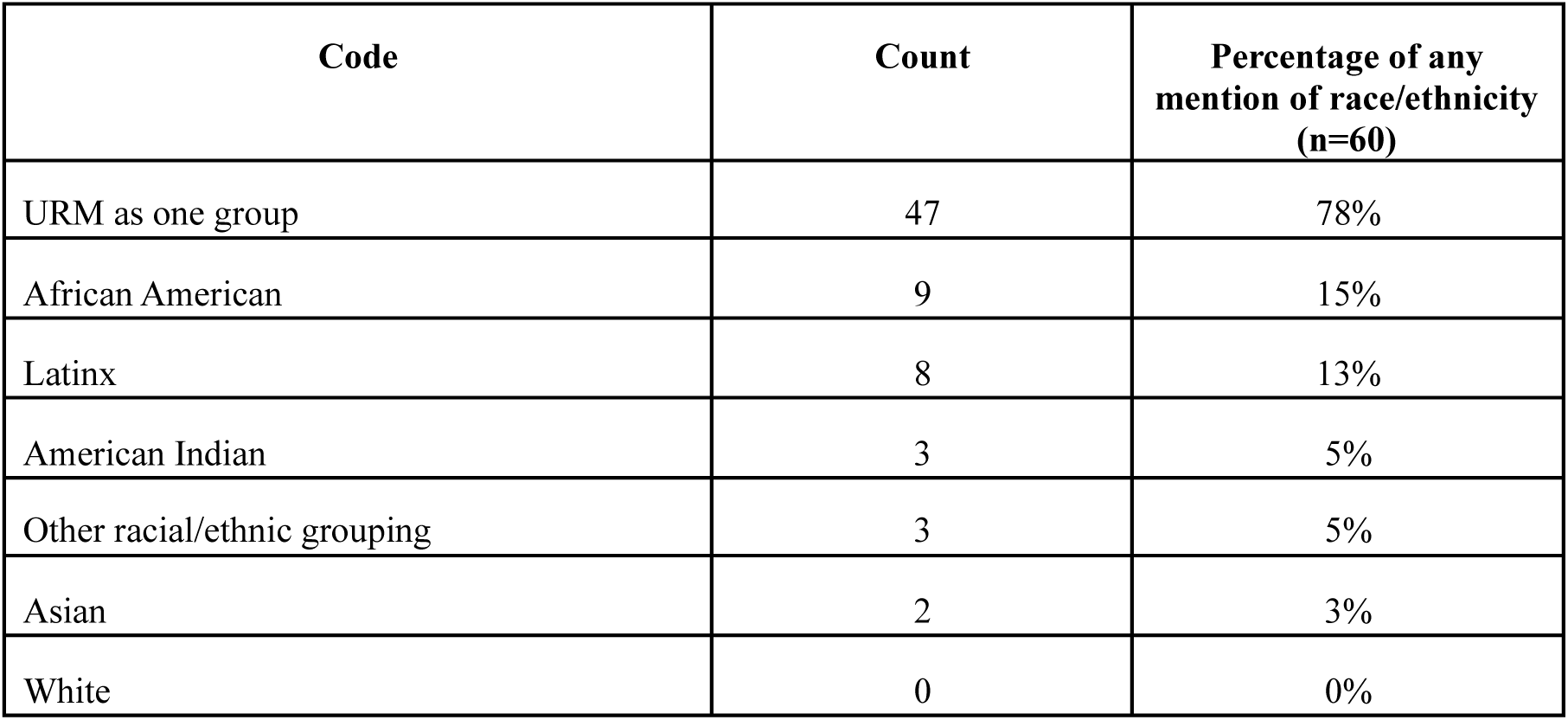
Number of abstracts mentioning particular racial/ethnic group emphasis

| Code | Count | Percentage of any mention of race/ethnicity (n=60) |
| --- | --- | --- |
| URM as one group | 47 | 78% |
| African American | 9 | 15% |
| Latinx | 8 | 13% |
| American Indian | 3 | 5% |
| Other racial/ethnic grouping | 3 | 5% |
| Asian | 2 | 3% |
| White | 0 | 0% |

Two of the 93 articles specified focusing on sexual and gender identities, with the two articles each focusing on both sexual *and* gender identities, lumping them together. The 91 remaining articles did not mention a focus on any sexual or gender identities.

Of the 93 articles, our coding determined that 92 of them articulated the use of at least one clearly specified research method. Most articles (n=68) described using one research method, while 19 identified two, and five identified three. Looking at the 92 articles together, noting that 24 used more than one method, Table 5 shows the distribution of methods across the articles. The most common methodological choices were quantitative: more than half of the abstracts described using survey methods, a quarter used grades, and a fifth school data.

**Table 5.** Number of abstracts using particular methods

| Code | Count | Percentage using method |
| --- | --- | --- |
| Survey | 43 | 58% |
| Grades | 24 | 26% |
| School Data | 18 | 20% |
| Interview | 16 | 17% |
| Observation | 9 | 10% |
| Other | 9 | 10% |
| Review of Literature | 2 | 2% |

Qualitative methods were less often used and included interviews (n=16) and observation (n=9).

Two articles employed reviews of the literature as their methods. Additionally, methods outside of those in our codebook were found in nine abstracts.

All of the 93 articles were able to be coded using one of the three codes developed by the team to reflect the level of intervention. However, one article was coded with two codes as it drew equally from both levels. A classroom level of analysis was the most common (n=37, 40%, see Table 6), followed by individual focus (n=30; 32%), and then structural approaches (n=27; 29%).

**Table 6.** Number of abstracts adopting particular levels of intervention

| Code | Count | Percentage using method |
| --- | --- | --- |
| Classroom | 37 | 39% |
| Individual | 30 | 32% |
| Structural | 27 | 29% |

## Discussion

To summarize our findings, research informing the biology education community draws heavily from psychological perspectives that are overwhelmingly not disaggregated (78% of articles identifying a group used a lumped together one), are by far more quantitative (58% used survey, 26% grades, 20% school data) than qualitative (17% used interview, 10% observation), and did not (72%) adopt structural approaches. Because the articles reviewed come from journals of national biology professional societies, they generally come from authors who have primary disciplinary training in the biological/life sciences. We speculate that training primarily in a biological discipline drives a bias for quantitative methods (over qualitative or mixed-method approaches), given their ubiquity and familiarity in these fields. This would also be true for administrators who move into positions of leadership from their academic units.

As the recognition of social factors in learning continues to meet with the priorities of institutions to use data to promote student retention, it is important to consider the range of perspectives available to inform decision making. BER would benefit from an expanded analytical and methodological repertoire. Including qualitative approaches (e.g., those from sociology and anthropology), ideally through interdisciplinary collaborations, could increase the ability to explain phenomena and support novel applications. This can furthermore help scholars take into fuller account the diversity of marginalized groups and look beyond deficit approaches.

A consideration of additional levels of analysis and an understanding of the various sociological and anthropological perspectives may therefore require additional professional development to build awareness and collaboration. While there is a great need for inclusion of qualitative and mixed-methods approaches in this work, we would recommend that rather than embark in independent training in a new way of thinking, and attempting to essentially take on the knowledge of a new field alone, there should be increased collaboration with those trained in qualitative methods, including sociologists, anthropologists, psychologists (and others) who likely teach and work with the same students on the same campus as biology educators.

Collaboration between those with a background in quantitative methods and biology, in combination with those trained in qualitative methods, will not only help biology education researchers to learn about research in this field, but likely strengthen the work of researchers from these varied disciplines.

In addition, our review of these articles showed that we did not find a large number of empirical articles on diversity, equity, and inclusion. In examining the aims and scopes of the national biology education society journals we investigated, we see that theoretical lenses and methodological practices are not emphasized, with focus instead on the application of the research to education in the biological sciences. Thus, we suggest that biology education research journals that want to impact diversity, equity, and inclusion, may want to consider expanding their aims, scope, or other criteria in order to draw on a wider and more diverse body of scholarship.

As shown, the biology education research in these journals draws heavily from psychological perspectives that are not disaggregated, are quantitative, and do not generally use structural approaches. Informed by broader changes in other STEM education research fields, it is increasingly clear that understanding the social aspects of higher education environments will be critical to broadening participation, particularly when it comes to Persons Excluded because of their Ethnicity or Race (PEERS, Asai, 2020; Fries-Britt & White-Lewis, 2020; Marshall et al., 2021; McGee, 2016).

There are some limitations of this work, many of which are balanced based on the goals we sought to achieve in this study. For example, analyzing only abstracts (rather than full articles) is a weakness, given that we are not able to make any inferences on the composition of samples in papers that did not specifically mention these concepts outside of their abstract.

Papers that did not emphasize disaggregating by race in their abstract, for example, might have actually done so in the body of the paper. Yet, identifying research methods and explanatory frame/level are more common attributes/expectations of research abstracts. We balanced the feasibility of sample size with the robust coding schema and methodologies we sought to use wherein we established a high degree of intercoder reliability, and benefitted from numerous coders and a larger team’s expertise in coming to consensus on that coding.

In addition, our work is limited by the search terms used. We note that the initial search terms used to gather articles for analysis are not exhaustive, and represent an informed and robust point of departure to make the claims we have made. Additional work using terms such as “marginalized” often favored by social scientists, especially more recently, would likely expand the pool. However, given that our search term creation was led by two biology education experts with decades of experience, we have confidence that they speak to the terms used by the community.

Our choice to include only the main biology education research journals created by societies eliminated BER work in other journals from being considered. For instance, we note that Dirks (Dirks, 2011) found a larger *quantity* of BER articles in DBER journals not included in this study, for example in the *Journal of Research in Science Teaching*, the *Journal of College Science Teaching*. Yet we wished to focus on journals particularly studying BER, and journals such as the *Journal of Research in Science Teaching* and the *Journal of College Science Teaching* will, by virtue of their broader STEM remit, have a different audience to BER-focused journals. This speaks back to the opening of the present manuscript as emphasizing the journal articles we did because of their influence on biology educators and researchers specifically.

In closing, we suggest that future research into this matter peel back additional layers to uncover more details about the specifical theoretical lenses (or lack thereof) deployed in biology education research. By understanding more about the extent to which such lenses, as well as the types of lenses that are utilized, those engaging in biology education research (often biologists) can engage in targeted training to develop these skills to advance inclusion. This could help transform the field of biology education and usher in research and consequential changes to advance inclusion in new and important ways.

## Acknowledgments

This material is based upon work supported by the National Science Foundation under Grant No. 2010716.

## Data Availability Statement

The data that support the findings of this study are available from the corresponding author upon request.

## Conflict of Interest Statement

The authors declare no conflict of interest.

## Notes

### Competing Interest Statement

The authors have declared no competing interest.

### Summary of Updates

Author affiliation and contact information correction.

## References

1. Abernethy, E. F., Arismendi, I., Boegehold, A. G., Colón-Gaud, C., Cover, M. R., Larson, E. I., Moody, E. K., Penaluna, B. E., Shogren, A. J., Webster, A. J., & Woller-Skar, M. M. (2020). Diverse, equitable, and inclusive scientific societies: Progress and opportunities in the Society for Freshwater Science. Freshwater Science, 39(3), 363–376. 10.1086/709129

2. Akman, O., Eaton, C. D., Hrozencik, D., Jenkins, K. P., & Thompson, K. V. (2020). Building Community-Based Approaches to Systemic Reform in Mathematical Biology Education. Bulletin of Mathematical Biology, 82(8), 109. 10.1007/s11538-020-00781-4

3. Aldridge, J. M., & McChesney, K. (2018). The relationships between school climate and adolescent mental health and wellbeing: A systematic literature review. International Journal of Educational Research, 88, 121–145. 10.1016/j.ijer.2018.01.012

4. American Society for Cell Biology. (n.d.). CBE--Life Sciences Education (LSE). Retrieved September 15, 2022, from https://www.lifescied.org/info-for-authors

5. American Society for Microbiology. (n.d.). About JMBE. Retrieved September 15, 2022, from https://journals.asm.org/journal/jmbe/about

6. Asai, D. J. (2020). Race Matters. Cell, 181(4), 754–757. 10.1016/j.cell.2020.03.044

7. Bandura, A. (1977). Social learning theory (p. 247). Prentice Hall.

8. Brownell, S. E., & Tanner, K. D. (2012). Barriers to faculty pedagogical change: lack of training, time, incentives, and…tensions with professional identity? CBE Life Sciences Education, 11(4), 339–346. 10.1187/cbe.12-09-0163

9. Bucholtz, M., Barnwell, B., Skapoulli, E., & Lee, J.-E. J. (2012). Itineraries of identity in undergraduate science. Anthropology & Education Quarterly, 43(2), 157–172. 10.1111/j.1548-1492.2012.01167.x

10. C. Bullock, E. (2017). Only STEM can save us? examining race, place, and STEM education as property. Educational Studies, 53(6), 628–641. 10.1080/00131946.2017.1369082

11. Campbell-Montalvo, R. A., Caporale, N., McDowell, G. S., Idlebird, C., Wiens, K. M., Jackson, K. M., Marcette, J. D., & Moore, M. E. (2020). Insights from the inclusive environments and metrics in biology education and research network: our experience organizing inclusive biology education research events. Journal of Microbiology & Biology Education, 21(1), 10–1128. 10.1128/jmbe.v21i1.2083

12. Campbell-Montalvo, R., Lucy Putwen, A., Hill, L., Metcalf, H. E., Sims, E. L., Peters, J. W., Zimmerman, A. N., Gillian-Daniel, D. L., Leibnitz, G. M., & Segarra, V. A. (2022a). Scientific Societies Integrating Gender and Ethnoracial Diversity Efforts: A First Meeting Report from Amplifying the Alliance to Catalyze Change for Equity in STEM Success (ACCESS+). Journal of Microbiology & Biology Education, 23(1). 10.1128/jmbe.00340-21

13. Campbell-Montalvo, R., Kersaint, G., Smith, C. A. S., Puccia, E., Skvoretz, J., Wao, H., Martin, J. P., MacDonald, G., & Lee, R. (2022b). How stereotypes and relationships influence women and underrepresented minority students’ fit in engineering. Journal of Research in Science Teaching, 59(4), 656–692. 10.1002/tea.21740

14. Campbell-Montalvo, R., Kersaint, G., Smith, C. A. S., Puccia, E., Sidorova, O., Cooke, H., Wao, H., Martin, J. P., Skvoretz, J., MacDonald, G., & Lee, R. (2022c). The influence of professional engineering organizations on women and underrepresented minority students’ fit. Frontiers in Education, 6. 10.3389/feduc.2021.755471

15. Campbell-Montalvo, R., Cooke, H., Smith, C. A. S., Puccia, E., Hughes Miller, M., Wao, H., and Skvoretz, J. (in press). Que(e)rying How Professional STEM Societies’ Serve Queer and Trans Engineering and Science Undergraduates. Educational Studies.

16. Cech, Erin A, & Blair-Loy, M. (2019). The changing career trajectories of new parents in STEM. Proceedings of the National Academy of Sciences of the United States of America, 116(10), 4182–4187. 10.1073/pnas.1810862116

17. Cech, E A, & Waidzunas, T. J. (2021). Systemic inequalities for LGBTQ professionals in STEM. Science Advances, 7(3). 10.1126/sciadv.abe0933

18. Cedillo, S. (2018). Beyond Inquiry: Towards the Specificity of Anti-Blackness Studies in STEM Education. *Canadian Journal of Science*, Mathematics and Technology Education, 18(3), 242–256. 10.1007/s42330-018-0025-0

19. Corwin, L. A., Morton, T. R., Demetriou, C., & Panter, A. T. (2020). A qualitative investigation of stem students’ switch to non-stem majors post-transfer. Journal of Women and Minorities in Science and Engineering, 26(3), 263–301. 10.1615/JWomenMinorScienEng.2020027736

20. Dirks, C. (2011). The Current Status and Future Direction of Biology Education Research. National Academies of Science, Engineering and Medicine. https://sites.nationalacademies.org/cs/groups/dbassesite/documents/webpage/dbasse_072582.pdf

21. Dolan, E. L. (2012). Biology education research--a cultural (R)evolution. CBE Life Sciences Education, 11(4), 333–334. 10.1187/cbe.12-09-0166

22. Dolan, E. L. (2015). Biology education research 2.0. CBE Life Sciences Education, 14(4), ed1. 10.1187/cbe.15-11-0229

23. Freeman, J. B. (2020). Measuring and resolving LGBTQ disparities in STEM. Policy Insights from the Behavioral and Brain Sciences, 7(2), 141–148.

24. Fries-Britt, S., & White-Lewis, D. (2020). In pursuit of meaningful relationships: how black males perceive faculty interactions in STEM. The Urban Review, 52(3), 521–540. 10.1007/s11256-020-00559-x

25. Garcia, N. M., López, N., & Vélez, V. N. (2018). QuantCrit: Rectifying quantitative methods through critical race theory. Race ethnicity and education, 21(2), 149–157. ttps://doi.org/10.1080/13613324.2017.1377675

26. Greenwood, R., Suddaby, R., & Hinings, C. R. (2002). Theorizing change: the role of professional associations in the transformation of institutionalized fields. Australasian Medical Journal, 45(1), 58–80. 10.5465/3069285

27. Her, T. (2019). The underrepresentation of Hmong American college students in STEM majors (Order No. 13903494). Available from ProQuest Dissertations & Theses Global. (2278078985). Retrieved from https://www.proquest.com/dissertations-theses/underrepresentation-hmong-american-college/docview/2278078985/se-2

28. Idsoe, E. M. C. (2016). The importance of social learning environment factors for affective well-being among students. Emotional and Behavioural Difficulties, 21(2), 155–166. 10.1080/13632752.2015.1053695

29. Intemann, K. (2009). Why diversity matters: understanding and applying the diversity component of the national science foundation’s broader impacts criterion. Social Epistemology, 23(3–4), 249–266. 10.1080/02691720903364134

30. International Union of Biochemistry and Molecular Biology. (n.d.). Biochemistry and Molecular Biology Education. Retrieved September 15, 2022, from https://iubmb.onlinelibrary.wiley.com/hub/journal/15393429/author-guidelines.html

31. Leibnitz, G. M., Gillian-Daniel, D. L., Greenler, R. M. C. C., Campbell-Montalvo, R., Metcalf, H., Segarra, V. A., Peters, J. W., Patton, S., Lucy-Putwen, A., & Sims, E. L. (2021). The inclusive professional framework for societies: changing mental models to promote diverse, equitable, and inclusive STEM systems change. Frontiers in Sociology, 6, 784399. 10.3389/fsoc.2021.784399

32. Leyva, L. A., McNeill, R. T., Balmer, B. R., Marshall, B. L., King, V. E., & Alley, Z. D. (2022). Black Queer Students’ Counter-Stories of Invisibility in Undergraduate STEM as a White, Cisheteropatriarchal Space. American Educational Research Journal, 000283122210964. 10.3102/00028312221096455

33. Leyva, L. A., McNeill, R. T., Marshall, B. L., & Guzmán, O. A. (2021). “It Seems like They Purposefully Try to Make as Many Kids Drop”: An Analysis of Logics and Mechanisms of Racial-Gendered Inequality in Introductory Mathematics Instruction. The Journal of Higher Education, 92(5), 784–814. 10.1080/00221546.2021.1879586

34. Leyva, L. A. (2016). An Intersectional Analysis of Latin@ College Women’s Counter-stories in Mathematics. Journal of Urban Mathematics Education. 10.21423/jume-v9i2a295

35. Lin, N. (2017). Building a network theory of social capital. In Social capital: theory and research (pp. 3–28). Routledge. 10.4324/9781315129457-1

36. Lo, S. M., Gardner, G. E., Reid, J., Napoleon-Fanis, V., Carroll, P., Smith, E., & Sato, B. K. (2019). Prevailing Questions and Methodologies in Biology Education Research: A Longitudinal Analysis of Research in CBE-Life Sciences Education and at the Society for the Advancement of Biology Education Research. CBE Life Sciences Education, 18(1), ar9. 10.1187/cbe.18-08-0164

37. Marshall, A., Pack, A. D., Owusu, S. A., Hultman, R., Drake, D., Rutaganira, F. U. N., Namwanje, M., Evans, C. S., Garza-Lopez, E., Lewis, S. C., Termini, C. M., AshShareef, S., Hicsasmaz, I., Taylor, B., McReynolds, M. R., Shuler, H., & Hinton, A. O. (2021). Responding and navigating racialized microaggressions in STEM. Pathogens and Disease, 79(5). 10.1093/femspd/ftab027

38. Martin, D. B., Groves Price, P., & Moore, R. (2019). Refusing systemic violence against black children. In J. Davis & C. C. Jett (Eds.), Critical race theory in mathematics education (pp. 32–55). Routledge. 10.4324/9781315121192-4

39. McGee, E. O., & Martin, D. B. (2011). “You would not believe what I have to go through to prove my intellectual value!” stereotype management among academically successful black Bmathematics and engineering students. American Educational Research Journal, 48(6), 1347–1389. 10.3102/0002831211423972

40. McGee, E. O. (2016). Devalued Black and Latino racial identities. American Educational Research Journal, 53(6), 1626–1662. 10.3102/0002831216676572

41. McGee, E. O., Thakore, B. K., & LaBlance, S. S. (2017). The burden of being “model”: Racialized experiences of Asian STEM college students. Journal of Diversity in Higher Education, 10(3), 253–270. 10.1037/dhe0000022

42. McGee, R. (2016). Biomedical workforce diversity: the context for mentoring to develop talents and foster success within the ‘pipeline’. AIDS and Behavior, 20 *Suppl 2*, 231–237. 10.1007/s10461-016-1486-7

43. McPherson, M., Smith-Lovin, L., & Cook, J. M. (2001). Birds of a feather: homophily in social networks. Annual Review of Sociology, 27(1), 415–444. 10.1146/annurev.soc.27.1.415

44. Mishra, S. (2020). Social networks, social capital, social support and academic success in higher education: A systematic review with a special focus on ‘underrepresented’ students. Educational Research Review, 29, 100307. 10.1016/j.edurev.2019.100307

45. Morton, Gee, & Woodson. (2020). Being vs. Becoming: Transcending STEM Identity Development through Afropessimism, Moving toward a Black X Consciousness in STEM. The Journal of Negro Education, 88(3), 327. 10.7709/jnegroeducation.88.3.0327

46. Morton, T. R., & Parsons, E. C. (2018). #BlackGirlMagic: The identity conceptualization of Black women in undergraduate STEM education. Science Education, 102(6), 1363–1393. 10.1002/sce.21477

47. Mourad, T. M., McNulty, A. F., Liwosz, D., Tice, K., Abbott, F., Williams, G. C., & Reynolds, J. A. (2018). The role of a professional society in broadening participation in science: A national model for increasing persistence. Bioscience, 68(9), 715–721. 10.1093/biosci/biy066

48. National Institutes of Health. (2017, December 6). NIH Policy and Guidelines on The Inclusion of Women and Minorities as Subjects in Clinical Research. https://grants.nih.gov/policy/inclusion/women-and-minorities/guidelines.htm

49. National Science Foundation. (2021). Women, Minorities, and Persons with Disabilities in Science and Engineering: 2021. Special Report NSF 21-321 (National Center for Science and Engineering Statistics, Trans.). National Science Foundation.

50. Nxumalo, F., & Gitari, W. (2021). Introduction to the Special Theme on Responding to Anti-Blackness in Science, Mathematics, Technology and STEM Education. *Canadian Journal of Science*, Mathematics and Technology Education. 10.1007/s42330-021-00160-8

51. Paris, Django. (2019) Naming beyond the white settler colonial gaze in educational research. International Journal of Qualitative Studies in Education, 32:3, 217–224. 10.1080/09518398.2019.1576943

52. Peffer, M., & Renken, M. (2016). Practical Strategies for Collaboration across Discipline-Based Education Research and the Learning Sciences. CBE Life Sciences Education, 15(4). 10.1187/cbe.15-12-0252

53. Pérez-Stable, E. J. (2018, June 27). Communicating the Value of Race and Ethnicity in Research. https://www.nih.gov/about-nih/what-we-do/science-health-public-trust/perspectives/sciencehealth-public-trust/communicating-value-race-ethnicity-research

54. Rosa, J., & Flores, N. (2017). Unsettling race and language: Toward a raciolinguistic perspective. Language in Society, 46(05), 621–647. 10.1017/S0047404517000562

55. Secules, S., Gupta, A., Elby, A., & Tanu, E. (2018). Supporting the narrative agency of a marginalized engineering student. Journal of Engineering Education, 107(2), 186–218. 10.1002/jee.20201

56. Secules, S., Sochacka, N. W., Huff, J. L., & Walther, J. (2021). The social construction of professional shame for undergraduate engineering students. Journal of Engineering Education. 10.1002/jee.20419

57. Segarra, V. A., Primus, C., Unguez, G. A., Edwards, A., Etson, C., Flores, S. C., Fry, C., Guillory, A. N., Ingram, S. L., Lawson, M., McGee, R., Paxson, S., Phelan, L., Suggs, K., Vega, L. R., Vuong, E., Havran, J. C., Leon, A., Burton, M. D., … Ramirez-Alvarado, M. (2020a). Scientific societies fostering inclusivity through speaker diversity in annual meeting programming: a call to action. Molecular Biology of the Cell, 31(23), 2495–2501. 10.1091/mbc.E20-06-0381

58. Segarra, V. A., Vega, L. R., Primus, C., Etson, C., Guillory, A. N., Edwards, A., Flores, S. C., Fry, C., Ingram, S. L., Lawson, M., McGee, R., Paxson, S., Phelan, L., Suggs, K., Vuong, E., Hammonds-Odie, L., Leibowitz, M. J., Zavala, M., Lujan, J. L., & Ramirez-Alvarado, M. (2020b). Scientific Societies Fostering Inclusive Scientific Environments through Travel Awards: Current Practices and Recommendations. CBE Life Sciences Education, 19(2), es3. 10.1187/cbe.19-11-0262

59. Shipley, T. F., McConnell, D., McNeal, K. S., Petcovic, H. L., & John, K. E. St. (2017). Transdisciplinary Science Education Research and Practice: Opportunities for GER in a Developing STEM Discipline-Based Education Research Alliance (DBER-A). Journal of Geoscience Education, 65(4), 354–362. 10.5408/1089-9995-65.4.354

60. Slaton, A. E., & Pawley, A. L. (2018). The power and politics of engineering education research design: saving the ‘small N.’ Engineering Studies, 10(2–3), 133–157. 10.1080/19378629.2018.1550785

61. Smith, Chrystal A S, Wao, H., Kersaint, G., Campbell-Montalvo, R., Gray-Ray, P., Puccia, E., Martin, J. P., Lee, R., Skvoretz, J., & MacDonald, G. (2021). Social capital from professional engineering organizations and the persistence of women and underrepresented minority undergraduates. Frontiers in Sociology, 6, 671856. 10.3389/fsoc.2021.671856

62. Smith, C A S, Wao, H., Martin, J., MacDonald, G. T., Lee, R., & Kersaint, G. (2015). Designing a Survey for Engineering Undergraduates using Free Listing -An Anthropological Structured Technique (Vol. 122).

63. Smith, J. L., Cech, E., Metz, A., Huntoon, M., & Moyer, C. (2014). Giving back or giving up: Native American student experiences in science and engineering. Cultural Diversity & Ethnic Minority Psychology, 20(3), 413–429. 10.1037/a0036945

64. Steele, C. M., & Aronson, J. (1995). Stereotype threat and the intellectual test performance of African Americans. Journal of Personality and Social Psychology, 69(5), 797–811. 10.1037/0022-3514.69.5.797

65. Tinto, V. (2006). Research and practice of student retention: what next? *Journal of College Student Retention: Research*, Theory & Practice, 8(1), 1–19. 10.2190/4YNU-4TMB-22DJ-AN4W

66. University of California Press. (n.d.). The American Biology Teacher. Retrieved September 15, 2022, from https://online.ucpress.edu/abt/pages/About

67. Vakil, S., & Ayers, R. (2019). The racial politics of STEM education in the USA: interrogations and explorations. Race Ethnicity and Education, 22(4), 449–458. 10.1080/13613324.2019.1592831

68. Wilton, M., Vargas, P., Prevost, L., Lo, S. M., Cooke, J. E., Gin, L. E., Imad, M., Tatapudy, S., & Sato, B. (2022). Moving towards More Diverse and Welcoming Conference Spaces: Data-Driven Perspectives from Biology Education Research Scholars. Journal of Microbiology & Biology Education, 23(2). 10.1128/jmbe.00048-22

69. Yonas, A., Sleeth, M., & Cotner, S. (2020). In a “scientist spotlight” intervention, diverse student identities matter. Journal of Microbiology & Biology Education, 21(1). 10.1128/jmbe.v21i1.2013

